# The Clustered Gamma Protocadherin Pcdhγc4 Isoform Regulates Cortical Interneuron Programmed Cell Death in the Mouse Cortex

**DOI:** 10.1101/2023.02.03.526887

**Authors:** Walter R Mancia Leon, David M Steffen, Fiona Dale-Huang, Benjamin Rakela, Arnar Breevoort, Ricardo Romero-Rodriguez, Andrea R Hasenstaub, Michael P Stryker, Joshua A Weiner, Arturo Alvarez-Buylla

## Abstract

Cortical function critically depends on inhibitory/excitatory balance. Cortical inhibitory interneurons (cINs) are born in the ventral forebrain and migrate into cortex, where their numbers are adjusted by programmed cell death. Previously, we showed that loss of clustered gamma protocadherins (*Pcdhγ*), but not of genes in the alpha or beta clusters, increased dramatically cIN BAX-dependent cell death in mice. Here we show that the sole deletion of the Pcdhγc4 isoform, but not of the other 21 isoforms in the Pcdhγ gene cluster, increased cIN cell death in mice during the normal period of programmed cell death. Viral expression of the *Pcdhγc4* isoform rescued transplanted cINs lacking *Pcdhγ* from cell death. We conclude that *Pcdhγ*, specifically *Pcdhγc4*, plays a critical role in regulating the survival of cINs during their normal period of cell death. This demonstrates a novel specificity in the role of *Pcdhγ* isoforms in cortical development.

## Introduction

In the cerebral cortex, inhibitory cortical neurons (cINs) are essential for sculpting, gating, and regulating neuronal excitation. Dysfunction or changes in the number of cINs is a hallmark of neurological disorders including epilepsy, schizophrenia, and autism (Chao et al., 2010; Lewis et al., 2005; Marín, 2012; Rossignol, 2011; Rubenstein & Merzenich, 2003; Verret et al., 2012). Ensuring the correct numbers of cINs are established during cerebral cortex development is an essential step in establishing proper brain function.

During embryonic development excess numbers of cINs are produced in the ventral forebrain within the medial and caudal ganglionic eminences (MGE and CGE). The young cINs then migrate dorsally into the cortex and form connections with locally produced excitatory neurons. Within the first two postnatal weeks of cortical development in mice, approximately 40% of cINs are eliminated by programmed cell death (Southwell et al., 2012; Wong et al., 2018). An intriguing feature of this developmental process of cell elimination is its timing and location. While most cINs are born in the ventral telencephalon between E11.5 and 16.5, their programmed cell death occurs postnatally (10-15 days later) and in the cortex, far from their birthplace. The timing of cIN programmed cell death correlates with several features of cortical development including the emergence of correlated activity and the development of cIN morphological complexity and synaptic connectivity (Ben-Ari et al., 2004; Connors et al., 1983; Priya et al., 2018; Seress et al., 1989; Tyzio et al., 1999; Yang et al., 2012). Indeed, changes in correlated neuronal activity have been associated with altered cIN survival. Increased correlated patterns of neuronal activity increased cIN survival while decreased cIN activity decreased cIN survival (Duan et al., 2020). Given the role of the clustered protocadherins in neuronal tiling, arborization, survival and axon targeting (Chen et al., 2012; Garrett et al., 2012, 2019; Katori et al., 2017; Lefebvre et al., 2008; Molumby et al., 2016; Mountoufaris et al., 2017; Prasad et al., 2008; Wang et al., 2002), we postulated that these diverse adhesion proteins could be required to establish initial cell-cell connectivity among neurons during early postnatal development and to regulate programmed cell death in the cortex.

The clustered protocadherins (Pcdhs) (Wu & Maniatis, 1999) are a set of fifty-eight cell surface homophilic-adhesion molecules that are tandemly arranged in three subclusters named alpha, beta, and gamma: *Pcdha, Pcdhb*, and *Pcdhg* (Wu et al., 2001). We have previously shown that the function of the isoforms in the *Pcdhg* cluster, but not those of the *Pcdha* or *Pcdhb*, are required for the survival of most cINs (Mancia Leon et al., 2020). Using transplantation to bypass neonatal lethality, we have also shown that the combined removal of three *Pcdhγ* isoforms (*Pcdhγc3, Pcdhγc4, and Pcdhγc5*) resulted in increased cell death. Within the *Pcdhg* cluster, the deletion of *Pcdhγc4*, but not that of other isoforms, is sufficient to cause neonatal lethality and increased cell death in the spinal cord (Garrett et al., 2019; Prasad et al., 2008). We wondered whether Pcdhγc4 also plays a unique and specific role in the regulation of cIN survival in the neocortex.

In the present study, we screened published single-cell RNA sequencing (scRNA-seq) datasets to determine whether *Pcdhγ*c4 is expressed in cINs in the adult mouse cortex. The *Pcdhγ*c4 was surprisingly enriched in the cIN population and largely absent from excitatory neurons in the adult cortex. In contrast, the *Pcdhγ*c5 isoform was mainly expressed in excitatory neurons (Tasic et al., 2018; Yao et al., 2021). Next, we characterized the expression of *Pcdhγ*c4 and *Pcdhγ*c5 during early postnatal development in the cortex of mice. Using RNAScope, we observed that the expression of *Pcdhγ*c5 was minimal at P5, increasing by P7. By P14, *Pcdhγ*c5 showed preferential expression in excitatory neurons. *Pcdhγ*c4 expression also increased from P5 to P7, and by P14 *Pcdhγc4* was found enriched in the MGE-derived cINs. Using a series of knockout mice in which various of the *Pcdhγ* isoforms were deleted (Garrett et al., 2019), combined with heterochronic transplantation (Mancia Leon et al., 2020), we then showed that the 19 A- and B-type *Pcdhγ* isoforms, as well as two Pcdhγ C-type isoforms (Pcdhγc3 and Pcdhγc5), have at most a minimal effect on cIN survival. In contrast, the single deletion of *Pcdhγ*c4 was sufficient to increase cell death dramatically among cINs derived from the MGE during the normal period of programmed cell death. Lastly, we showed that *Pcdhγ*-deficient cINs were rescued from excess cell death by the viral over-expression of the *Pcdhγ*c4 isoform. We conclude that *Pcdhγ* diversity is not required for cIN survival; rather the expression of a single isoform, that of *Pcdhγ*c4, is necessary and sufficient for the survival of most cINs.

## Results

### Pcdhγ C-type expression in the mouse cortex during programmed cell death

The *Pcdhγ* gene locus encodes 22 distinct Pcdhγ proteins, which are subclassified as A-, B-, or C-type isoforms. Previous work suggests that *Pcdhγ* and specifically the C-type isoforms play a key role in the regulation of cIN programmed cell death (Mancia Leon et al., 2020). This previous work suggested that either individual or multiple C-type *γ*-Pcdhs are required to maintain appropriate numbers of cINs in the cortex. In order to determine whether differential expression patterns of individual C-type *γ-*Pcdhs might be involved in cIN survival, we screened scRNA sequencing datasets to determine which C-type *γ*-Pcdhs are expressed in cIN. Interestingly, two scRNA sequencing datasets generated from adult animals (>P50) (Tasic et al., 2018; Yao et al., 2021) showed a distinct increase in the expression of *Pcdhγc4* in GABAergic cells. In contrast, cells identified as glutamatergic excitatory neurons expressed higher levels of *Pcdhγc*5 **(Fig. 1A-B)**. We next looked at the expression of *Pcdhγc4* and *Pcdhγc5* in the different sub-classes of both excitatory and inhibitory neurons **(Fig.1C-D)**. Among inhibitory populations, interferon gamma-induced GTPase (Igtp), Neuron-derived neurotrophic factor (Ndnf), Parvalalbumin (Pvalb), Somatostatin (SST), and Vasoactive intestinal peptide-expressing (Vip) subtypes showed a higher expression of *Pcdhγc4* and lower levels of *Pcdhγc5*. Among the excitatory neurons, higher levels of *Pcdhγc5* expression was especially evident among deep and superficial glutamatergic neurons. Together, these scRNA-seq datasets indicate that the *Pcdhγc4* and *Pcdhγc5* genes are differentially expressed between cINs and excitatory neurons in the adult mouse cortex. Interestingly, expression of *Pcdhγc3*, which plays a unique role in promoting cortical dendrite arborization through its signaling partner Axin1 (Steffen et al., 2023), showed no distinct expression bias between GABAergic and Glutamatergic cells and was expressed at low levels similar to that of the other *Pcdhγ* genes **(Fig. 1A-B)**.

**Figure 1.**
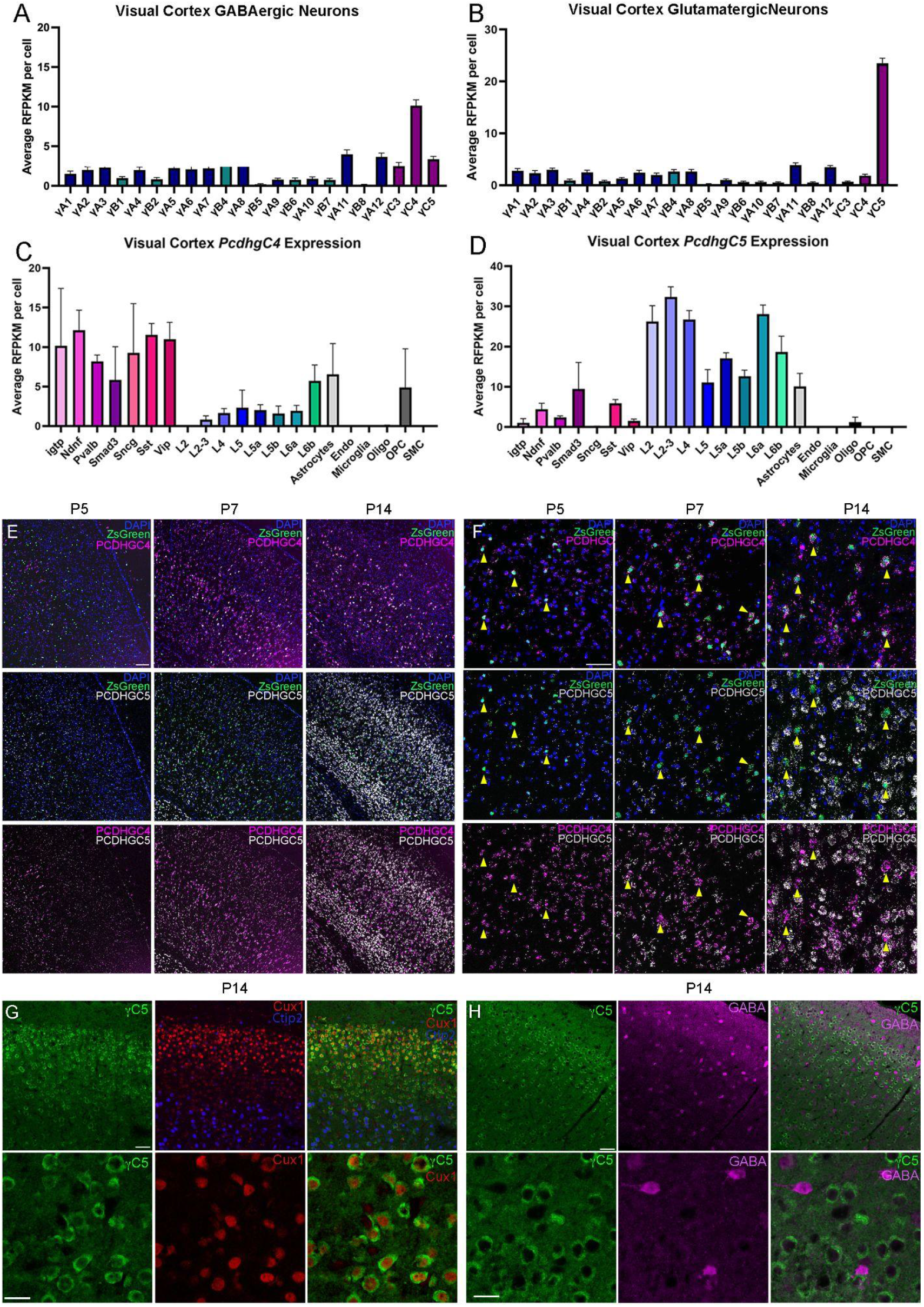
Pcdhγ C-type expression in the mouse cortex during programmed cell death. **A-B.** ScRNA-seq from previously generated dataset show expression bias of *Pcdhγc4* in GABAergic cIN (A), and *Pcdhγc5* in Glutamatergic neurons (B).Error bars represent the error of the mean. **C-D.** Expression levels of *Pcdhγc4* is higher in inhibitory neuron subtypes (C), while *Pcdhγc5* expression is higher in Glutamatergic neuron subtypes (D). Error bars represent the error of the mean. interferon gamma-induced GTPase (Igtp), Neuron-derived neurotrophic factor (Ndnf), Parvalalbumin (Pvalb), Somatostatin (SST), and Vasoactive intestinal peptide-expressing (Vip). **E-F.** RNA scope of Nkx2.1;Ai6 cortex during programmed cell death. Low magnification (E) and higher magnification (F) shows expression of *Pcdhγc4* and *Pcdhγc5* to be minimal at *Pcdhγc5*, *Pcdhγc4* expression is increased by P7 and is expressed in MGE-derived ZsGreen+ cells (yellow arrows). At P14, expression of both *Pcdhγc4* and *Pcdhγc5* is high, but distinctly not in the same cells, with *Pcdhγc4* being highly expressed in ZsGreen+ MGE-derived cINs. Scale bar low-magnification (G) = 100 um. Scale bar high-magnification (F) = 50 um. **G-H.** IHC in P14 mouse cortex shows Pcdhγc5 association with excitatory neurons (Ctip2 and Cux1)(G), and not with GABAergic cIN (H). Scale bar top row = 50 um. Scale bar bottom row = 25 um.

In order to corroborate the above expression pattern histologically, we performed *in-situ hybridizations* using RNAscope, in combination with a reporter mouse in which MGE-derived cINs were labeled with a ZsGreen protein (Nkx2.1;Ai6 mouse line). Using validated probes for the *Pcdhγc4* and *Pcdhγc5* mRNA transcripts, we looked at multiple times during the period of programmed cell death (P5, P7, and P14). At P5, we found low expression of *Pcdhγc4* and *Pcdhγc5* in both ZsGreen+ or - cells. However, a subpopulation of ZsGreen+, MGE-derived cINs at P5 was associated with a higher *Pcdhγc4* signal **(Fig. 1E-F)**. By P7, both *Pcdhγc4 and Pcdhγc5* were observed in the majority of cells, although the *Pcdhγc4* signal associated with the ZsGreen+ neurons was more robust **(Fig. 1E-F)**. At P14, when the period of programmed cIN cell death is largely over, we found a dramatic increase in *Pcdhγc4* and *Pcdhγc5* expression, consistent with previous reports of Pcdh*γ*c5 expression beginning in the second postnatal week in rat cortex (Li et al., 2010). Cortical neurons at this time showed a clear preferential expression of either the *Pcdhγc4* or the *Pcdhγc5* isoform. ZsGreen+ neurons showed preferential expression of the *Pcdhγc4* isoform, while many of the ZsGreen-cells showed lower or no expression of *Pcdhγc4*. Next, we aimed to validate these findings using IHC. While we were unable to procure a reliable antibody for Pcdhγc4, we did find an antibody that specifically labels the Pcdhγc5 isoform (validated using 3R2 mice lacking *Pcdhγc5* gene; Garrett et al., 2019; data not shown). We co-stained for Pcdhγc5 and markers for upper-layer excitatory neurons (Cux1) and lower-layer excitatory neurons (Ctip2). At P14, Pcdhγc5 expression clearly surrounded neuronal nuclei expressing these excitatory transcription factor markers **(Fig. 1G)**. To determine if Pcdh*γ*c5 expression is lower in inhibitory neurons, we also stained P14 cortex for Pcdh*γ*c5 and GABA **(Fig. 1H)**. Consistent with the scRNA-seq data, GABA-expressing inhibitory neurons expressed low/background levels of Pcdhγc5. Together, these data indicate divergent expression of *Pcdhγc4* isoforms in cINs and *Pcdhγc5* isoforms in excitatory neurons, which is a pattern that emerges during the period of programmed cell death and is maintained in the adult cortex.

### Pcdhγc4 deletion results in the elimination of most cINs

Our previous work and that of others has shown that the combined deletion of the three *Pcdhγ* C-type genes (*Pcdhγc3, Pcdhγc4*, and *Pcdhγc5*) results in increased cIN cell death during the period of naturally occurring cell death (Chen et al., 2012; Mancia Leon et al., 2020). Other studies have shown that constitutive genetic deletion of *Pcdhγc4*, but not that of *Pcdhγc3 or Pcdhγc5*, leads to neonatal lethality and increased interneuron cell death in the spinal cord (Garrett et al., 2019). To ascertain the function of *Pcdhγc4* in the regulation of cIN programmed cell death, we used mice with constitutive deletions of the *Pcdhγc4* isoform (Pcdhγ^C4KO^ mice; Garrett et al., 2019). Pcdhγ^C4KO^ mice were crossed to the Nkx2.1^Cre^; Ai14 MGE/preoptic area (POA) reporter mouse line to provide a genetically encoded fluorescent label for the cINs (**Fig. 2A**).

**Figure 2.**
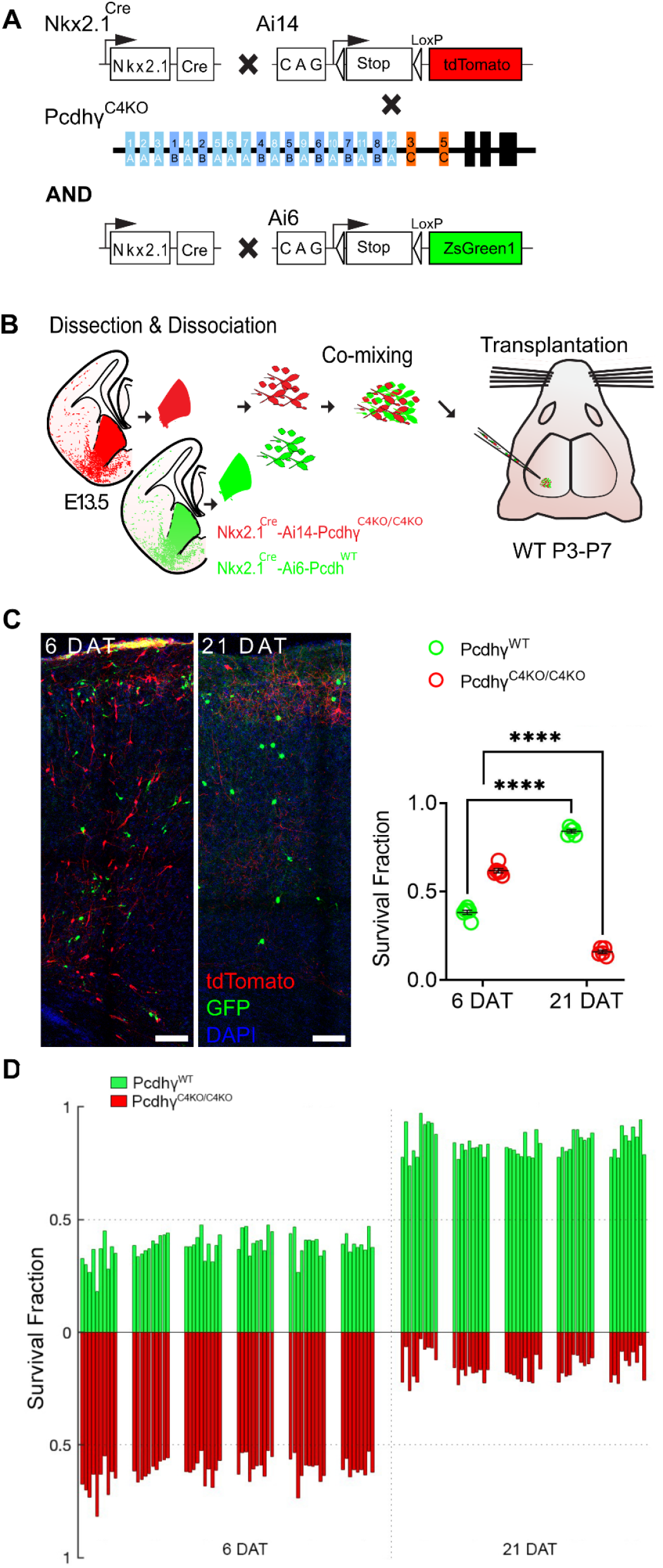
Genetic deletion of Pcdhγc4 increased cell death in MGE-derived cINs. **A.** Diagram of genetic crosses between MGE/POA-specific reporter and Pcdhγ^C4KO^ mice. Pcdhγ^C4KO^ homozygous MGE cells were obtained from the *Nkx2.1^Cre^*; Ai14;Pcdhγ^C4KO/C4KO^ embryos, whereas control cells were obtained from *Nkx2.1^Cre^*; Ai6 embryos. **B.** Schematics of transplantation protocol. The MGEs from E13.5 Pcdhγ^C4KO^ homozygous mutant or control embryos were dissected, dissociated, and mixed in similar proportions. The mixture of GFP+ (Pcdhγ^WT^) and tdTomato+ (Pcdhγ^C4KO/C4KO^) cells was grafted into the cortex of WT neonate mice. **C.** Left - Confocal images from the cortex of 6 and 21 DAT mice. The transplanted cells are labeled with GFP (Pcdhγ^WT^) or tdTomato (Pcdhγ^C4KO/C4KO^). Right - Quantifications (shown as survival fraction) of surviving MGE-derived cINs at 6 and 21 DAT. Both the transplanted GFP and tdTomato-labeled cells undergo programmed cell death between 6 and 21 DAT, but the Pcdhγ^C4KO/C4KO^ cells are eliminated at significantly higher rates. **D.** Survival fraction quantification from (C) shown by the brain section (each bar) and separated by animals at 6 and 21 DAT. Scale bar = 50 um, Nested-ANOVA, ****p =3.147e-10, n = 5 mice per time point and 10 brain sections quantified per mouse

Embryos homozygous for deletion of the *Pcdhγc4* isoform develop normally with no apparent weight, size or brain abnormalities and are born in normal Mendelian ratios. However, these mice die perinatally (within 24 hours of birth) before the period of programmed cell death for cINs around P7. In order to bypass lethality and to study the role of Pcdhγc4 deletion in cIN survival postnatally, we utilized a co-transplantation method of cINs that are WT or mutant (**Fig. 2B**). This allowed us to compare the survival of mutant and WT cells within the same WT environment. The F2 generation of Nkx2.1^Cre^;Ai14;Pcdhγ^C4KO/+^ mice were bred to generate E13.5 embryos, homozygous for the *Pcdhγ^C4KO^* allele. In these embryos, MGE/POA-derived cells lacking Pcdhγc4 are fluorescently labeled with the tdTomato protein upon Cre-driven recombination of the Ai14 allele.

The MGE of embryos carrying the homozygous deletion of the Pcdhγc4 allele was microdissected. As a control, GFP-expressing cINs were derived from microdissected MGEs of Nkx2.1^Cre^;Ai6, in which MGE-POA-derived cells are fluorescently labeled with ZsGreen (Madisen et al., 2010) (**Fig. 2A & B**) or Gad67-GFP embryos (**Supplementary Fig. 2A & B**). The MGEs were dissociated and cells were mixed in similar proportions (GFP and tdTomato). The mixture of GFP cells with tdTomato cells was transplanted into the cortex of host neonatal mice (P3-P7). The homozygous Pcdhγ^C4KO/C4KO^ cells were identified via tdTomato expression while cells carrying the Pcdhγ^WT^ allele were identified via expression of GFP (**Fig. 2C**). Survival of the transplanted cells was analyzed at 6 and 21 days post-transplant (DAT), which corresponds to postnatal days (P) 0 and 15 for the transplanted cells, cellular ages that span the period of endogenous cIN programmed cell death. At 6 DAT, the proportion of the fluorescently labeled transplanted cells expressing GFP (Pcdhγ^WT^) was 38.2%, and the tdTomato-positive Pcdhγ^C4KO/C4KO^ cells made up the balance, 61.8%, of all transplanted cells that survived. Importantly, there was no apparent change in the proportion of GFP to tdTomato cells between 4 and 6 DAT (data not shown), before the period of naturally occurring cell death for the transplanted cells. By 21 DAT, however, the proportion of GFP to tdTomato cells had shifted dramatically, indicating that one cell population was eliminated at higher rates. Indeed, the tdTomato-positive Pcdhγ^C4KO/C4KO^ cell population dropped from 61.8 % (at 6 DAT) to 15.9% at 21 DAT. Conversely, the proportion of GFP-positive Pcdhγ^WT^ cINs had increased from 38.2% (at 6 DAT) to 84.1% at 21 DAT. Note that the increase in the proportion of GFP-positive cells does not reflect an increase in survival, but rather that the tdTomato labeled cells were eliminated in much greater numbers during programmed cell death. Importantly, similar results were found in experiments where we co-transplanted Pcdhγ^C4KO/C4KO^ with Pcdhγ^WT^ Cells derived from Gad67-GFP embryos (**Supplementary Fig. 2**). These results suggest that cells that lack the Pcdhγc4 protein are more likely to be eliminated during the normal period of cell death than those cells that express the WT Pcdhγc4.

### Deletion of all Pcdhγ, except Pcdhγc4, is sufficient for the survival of the majority of cINs

We wondered whether Pcdhγc3 and Pcdhγc5 may also contribute to cIN survival. In our previous study, deletion of the C-type isoforms (*Pcdhγc3, Pcdhγc4, and Pcdhγc5*) resulted in the elimination of most cINs, to levels similar to those observed after the loss of function of the entire Pcdhγ cluster (Mancia Leon et al., 2020). These suggested that the 19 alternate A and B-type Pcdhγ isoforms do not significantly contribute to the survival of cINs. Here, we used the Pcdhγ^1R1^ mouse line (Garrett et al., 2019), which lacks all 19 A and B-type *Pcdhγ* isoforms, as well as *Pcdhγc3* and *Pcdhγc5*, but retains *Pcdhγc4* (**Fig. 3A)**.

**Figure 3.**
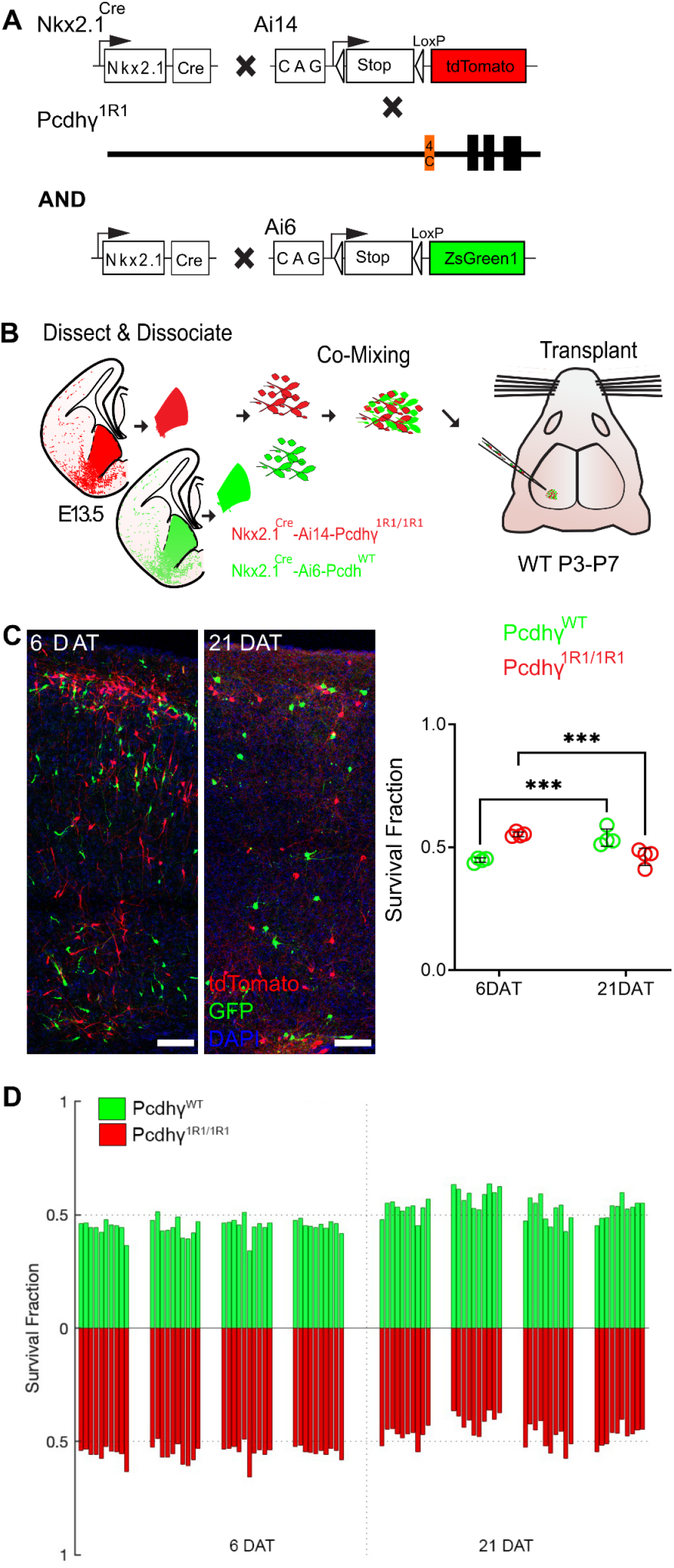
Most MGE-derived cINs survive the deletion of most Pcdhγ, except for Pcdhγc4. **A.** Diagram of genetic crosses between MGE/POA-specific reporter *Nkx2.1^Cre^*;Ai14 and Pcdhγ^1R1^ mice. Control cells were derived from *Nkx2.1^Cre^;Ai6* mice while Pcdhγ^1R1^ mutant cells were derived from *Nkx2.1^Cre^;Ai14*; Pcdhγ^1R1/1R1^ mice. **B.** Diagram of the transplantation protocol. The MGEs from E13.5 Pcdhγ^1R1^ homozygous or control embryos were dissected, dissociated, and mixed in similar proportions. The mixture of GFP+ (Pcdhγ^WT^) and tdTomato+ (Pcdhγ^1R1/1R1^) cells was grafted into the cortex of WT neonate mice. **C.** Left - Confocal images from the cortex of 6 and 21 DAT mice. The transplanted cells are labeled with GFP (Pcdhγ^WT^) or tdTomato (Pcdhγ^1R1/1R1^). Right - Quantifications of the transplanted cells that survived at 6 and 21 DAT. Note that both the GFP or tdTomato labeled cells underwent programmed cell death between 6 and 21 DAT, but both cell types survived to similar levels. **D.** Survival fraction quantification from (C) shown by the brain section (each bar) and separated by animals at 6 and 21 DAT. Scale bar = 50 um, Nested-ANOVA, ***p = 0.0026, n = 4 mice per time point and 10 brain sections per mouse.

Mice homozygous for the Pcdhγ^1R1^ allele (Pcdhγ^1R1/1R1^) are born in normal mendelian ratios, develop normally and are fertile, as previously reported (Garrett et al., 2019). Pcdhγ^1R1/R1^ mice were crossed to the above Nkx2.1^Cre^; Ai14 mouse line to label MGE/POA-derived cINs with the tdTomato protein (**Fig. 3A)**. As above, we used the co-transplantation to compare the survival of cINs solely expressing Pcdhγc4 to that of cINs expressing all 22 Pcdhγ isoforms. cIN precursor cells homozygous for the Pcdhγ^1R1/R1^ allele were obtained from E13.5 Nkx2.1^Cre^;Ai14;Pcdhγ^1R1/R1^ embryos. As a control, we used cIN precursor cells expressing WT Pcdhγ obtained from Nkx2.1^Cre^;Ai6 embryos (**Fig. 3B)**. Survival of the transplanted cells was analyzed at 6 and 21 DAT and is represented as the proportion of GFP to tdTomato cells at these time points. At 6 DAT, roughly 44.7% of all transplanted cells were GFP-positive (WT Pcdhγ) while the remaining cells (55.2%) were tdTomato positive (Pcdhγ^1R1/R1^). As hypothiszed, the survival of the Pcdhγ^1R1/1R1^ tdTomato positive cells was remarkably similar to that of the GFP-labeled control cells **(Fig. 3C-D)**. However, there was a small but significant drop in the Pcdhγ^1R1/1R1^ population (9.1%), as compared to a 45.9% decrease in the Pcdhγ^C4KO/C4KO^ transplants **(Fig. 2C-D)**. Similar results were observed when control cINs cells were obtained from the Gad67-GFP mouse line (**Supplementary Fig. 3**). Importantly, the proportion of GFP to tdTomato positive cells remained unchanged between 4 to 6 DAT (data not shown). Together, these data suggest that expression of Pcdhγc3 or Pcdhγc5 is not required for the survival of most cINs. Furthermore, these observations indicate that A- and B-type Pcdhγ isoforms are also not required for cIN survival. Given the small drop in cIN survival in Pcdhγ^1R1/R1^ cIN, we cannot completely rule out the possibility that Pcdhγc3 or Pcdhγc5 may make a small contribution to cIN survival. However, Pcdhγ^1R1/R1^ mice have been shown to express significantly decreased levels of Pcdhγc4 compared to WT mice (Garrett et al., 2019), which might explain the observed small, but statistically significant drop in the survival of the Pcdhγ^1R1/1R1^ cINs.

### Exogenous expression of Pcdhγc4 in Pcdhγ knockout cINs rescues cINs from cell death

The results above suggest that most cINs lacking expression of Pcdhγc4 are eliminated during the normal period of programmed cell death; in contrast, cINs expressing Pcdhγc4 as their only Pcdhγ isoform survive at nearly normal levels. We next asked whether cINs lacking expression of all twenty-two Pcdhγ isoforms could be rescued from cell death by re-introducing the Pcdhγc4 isoform. Since cINs lacking the function of all Pcdhγ isoforms are nearly eliminated when co-transplanted with WT cINs expressing Pcdhγ (**see figure 10E in (Mancia Leon et al.,** 2020), we reasoned that a rescue effect would be most evident in these mutant cells.

Dissociated MGE cells from E13.5 of Nkx2.1^Cre^;Ai14;Pcdhγ^fcon3/fcon3^ embryos were infected in suspension with lentivirus expressing the Pcdhγc4 isoform fused to GFP (**Fig. 4A & B)**. The infected cells were transplanted into WT host neonate mice and survival was analyzed at 6 and 21 DAT. The transplanted cINs expressing lentivirus-driven Pcdhγc4 were identified via the co-expression of tdTomato and GFP (**Fig. 4C**). Pcdhγ mutant cells that expressed only tdTomato were identified as not infected. We compared the survival of *Pcdhγc4*-transduced (GFP+tdTomato+) to non-transduced (tdTomato+ GFP-) at 6 and 21 DAT. While both the *Pcdhγc4*-transduced and non-transduced cINs undergo a wave of cell death between the above time points, the fraction of tdTomato+GFP+ cells increased between 6 and 21 DAT (33% to 68%). In contrast, the fraction of tdTomato+GFP-cells decreased from 66% to 32% during the same period of time (**Fig. 4 C)**. To control for possible non-specific effects of viral infection on cIN survival, we infected Nkx2.1^Cre^;Ai14 MGE cells that carry the WT Pcdhγ allele with lentivirus expressing GFP, and analyzed their survival at 6 and 21 DAT (**Supplementary Fig. 4A & B**). As above, we compared the survival of *Pcdhγc4*-transduced (tdTomato+GFP+) to non-transduced (tdTomato+GFP-) cells. At 6 DAT, the fraction of infected cells was approximately 50% and remained nearly constant between 6 and 21 DAT, indicating that infection with lentivirus and expression of GFP had no impact on cIN survival (**Supplementary Fig. 4C & D**). These results indicate that the introduction of Pcdhγc4 to cINs lacking endogenous Pcdhγ genes is sufficient to rescue the mutant cells from undergoing excessive apoptosis.

**Figure 4.**
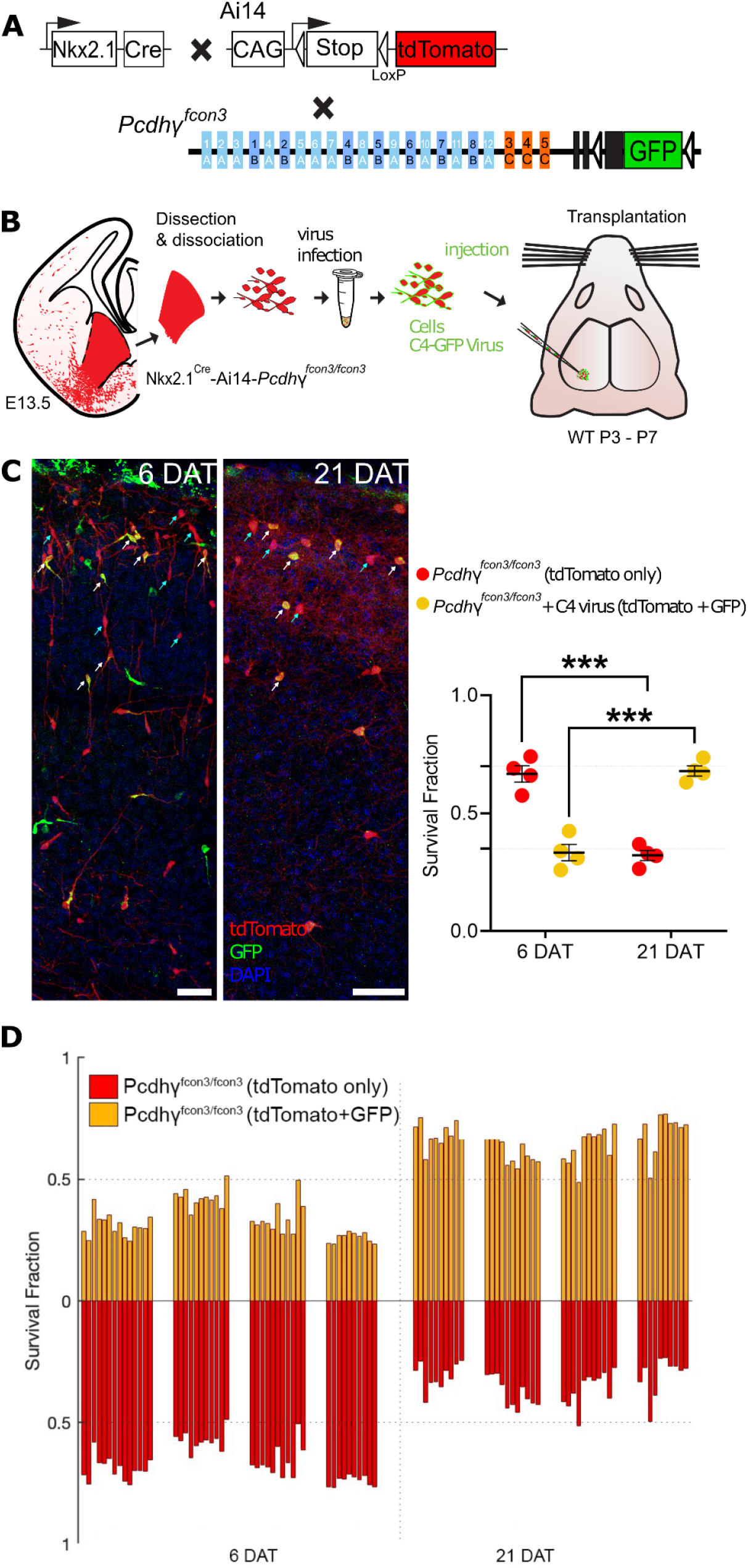
Lentiviral expression of Pcdhγc4 rescues cINs with Pcdhγ loss of function. **A.** Diagram of mouse genetic crosses. Pcdhγ^fcon3^ mice were crossed to the *Nkx2.1^Cre^;Ai14* mouse line to generate embryos with loss of function of all Pcdhγ isoforms. **B.** Schematic of the lentiviral infection and transplantation of MGE cIN precursors. The MGEs of *Nkx2.1^Cre^;Ai14;Pcdhγ^fcon3/fcon3^* embryos were dissected, dissociated, and infected in suspension with lentivirus carrying *Pcdhγc4-GFP*. The mixture of transduced and non-transduced cells was grafted into the cortex of WT neonate recipient mice. **C.** Confocal images of the transplanted cINs in the cortex at 6 and 21 DAT. Notice the expression of GFP can be found near the cell surface including the cell processes, reflecting the putative location of the transduced Pcdhγc4-GFP protein. Quantifications of the tdTomato+GFP- (teal arrows) or tdTomato+GFP+ (yellow cells, white arrows) cells are shown as the fraction of cells from the total tdTomato+ cells at 6 and 21 DAT. The fraction of the Pcdhγc4-transduced yellow cells increases from 6 to 21DAT, while the fraction of non-transduced (tdTomato+ GFP-) decreases at equivalent time points. **D.** Survival fraction quantification from (C) shown by the brain section (each bar) and separated by animals at 6 and 21 DAT. Scale bar = 50 um, Nested-ANOVA, ****p = 0.0002, n = 4 mice per time point and 10 brain sections per mouse from one transplant cohort.

## Discussion

This study reveals the preferential expression of Pcdhγc4 in MGE-derived cINs, in contrast to excitatory neurons, which preferentially express the distinct Pcdhγc5 isoform. This segregated pattern of expression develops during the period of programmed cell death, which also coincides with other key maturational stages. Soon after migration, cINs must integrate into cortical circuits, forming nascent synapses with both inhibitory and excitatory neurons. During this period local inhibitory networks require increased interneuron-interneuron connectivity to facilitate synchronous activity which promotes interneuron survival (Duan et al., 2020). Opposing expression of Pcdhγc5 and Pcdhγc4 on excitatory and inhibitory neuron adhesion interfaces, respectively, could help segregate the two populations of neurons and be a key mechanism in the formation of initial cortical circuits.

Previous studies show that Pcdhγ C-type isoforms are required to regulate neuronal numbers during the critical window of programmed cell death (Chen et al., 2012; Mancia Leon et al., 2020). The timing of Pcdhγc4 enrichment during postnatal development suggests that this specific Pcdhγ isoform could be playing a key role in programmed cell death. Indeed a recent study of the spinal cord and retina reported that the deletion of Pcdhγc4, but not that of other Pcdhγ isoforms, leads to increased cell death and neonatal lethality (Garrett et al., 2019). However, cIN-programmed cell death occurs mostly postnatally, whereas cell death in the spinal cord takes place prenatally (Prasad et al., 2008; Wang et al., 2002). Since mice lacking Pcdhγc*4 (Pcdhγ^C4KO^*) die soon after birth, we could not study the extent of cIN cell death directly in these mutant animals. We took advantage, therefore, of transplantation strategies to compare the survival of cells lacking *Pcdhγc4* to cells that carry all Pcdhγ isoforms. Pcdhγ^C4KO^ mice lack expression of *Pcdhγc4* but express the 19 A and B type Pcdhγ isoforms as well as *Pcdhγc3* and *Pcdhγc5*. Here, we show that most Pcdhγ^C4KO^ cINs are eliminated during the period of programmed cell death, a result that is consistent with that found with the deletion of the entire *Pcdhγ* gene cluster or of the three *Pcdhγ* C-type isoforms (Carriere et al., 2020; Chen et al., 2012; Mancia Leon et al., 2020). The above finding also suggests that C-type isoforms Pcdhγc3 or Pcdhγc5 do not play a major role in the regulation of cIN survival; this is consistent with the relatively low expression of these isoforms seen in adult cINs. Indeed, when we studied the survival of cINs that solely express Pcdhγc4 using cells from Pcdhγ^1R1^ embryos, we found that the majority of cINs survived at levels similar to those of cINs that carry all Pcdhγ isoforms. However, transplanted Pcdhγ^1R1^ cINs did show a small but significant reduction in survival, compared to control cells. While we cannot completely exclude a role for other Pcdhγ isoforms, including Pcdhγc3 or Pcdhγc5 in the Pcdhγ^1R1^ cells, the observed reduction in survival is likely due to reduced levels of Pcdhγc4 expression (~10% of WT levels) in Pcdhγ^1R1^ mice (Garrett et al., 2019). Indeed, this low expression of *Pcdhgc4* in Pcdhγ^1R1^ mice makes the survival of their cINs a more remarkable result. In order to test directly the function of Pcdhγc4, we used lentiviral driven expression of Pcdhγc4 in cINs that lack the function of all 22 Pcdhγ isoforms. Expression of Pcdhγc4 resulted in a dramatic increase in the survival of MGE-derived cINs when compared to cells lacking the function of the entire Pcdhγ cluster.

Our findings above suggest that Pcdhγ diversity is not required for the survival of cINs and show a unique and specific role for Pcdhγc4 in regulating the survival of cINs. Together, our results align with findings in the spinal cord and retina (Chen et al., 2012; Garrett et al., 2019), suggesting a potential general Pcdhγc4-driven mechanism that regulates the survival of local-circuit neurons across the CNS. While our data suggest that Pcdhγ diversity is dispensable for cIN survival, the diversity requirements for clustered Pcdhs in the *Pcdha* and *Pcdhb* clusters was not studied here: in all genotypes examined, these clusters remained intact. A recent study found that mutant mice lacking all call *Pcdha, Pcdhb*, and A- and B-type *Pcdhg* genes (thus retaining only Pcdhγc3, Pcdhγc4, and Pcdhγc5), also exhibited neonatal lethality and increased neuronal apoptosis (Kobayashi et al., 2023). These results are consistent with previous reports showing that Pcdhγc4 (uniquely among Pcdhγ isoforms) requires other “carrier” cPcdhs to be transported to the cell-surface (Thu et al. 2014). While the mutant mice used by Kobayahsi et al. (2022) retain the Pcdh*γ*c3 and Pcdh*γ*c5 isoforms, these are unlikely to act as sufficient carriers due to low embryonic expression of Pcdhγc3 and Pcdhγc5 (Kobayashi et al. 2022) and low intra-family affinity for *cis*-interactions required for carriers (Goodman et al., 2022). During normal development, it is also possible that Pcdhγc4 forms heterodimers, *in cis*, with members of *Pcdha* and/or *Pcdhb* clusters, providing cellular specificity that is important for cIN survival. In both scenarios, our findings suggest a unique role in which Pcdhγc4 regulates programmed cell death. However, the mechanisms by which the Pcdhγc4 isoform regulates apoptosis among young cINs, leading to the adjustment of local circuit neuron numbers, remains unknown. Presumably, this isoform has unique intracellular signaling partners; precedents for this possibility comes from a recent study showing that Pcdh*γ*C3 plays a unique role in promoting cortical pyramidal neuron dendritic arborization through intracellular interactions with Axin1 (Steffen et al., 2023).

Together this work, along with previous studies, suggests that the final number of cINs in the cerebral cortex is in part determined through a cell-or population-intrinsic mechanism involving cell-cell interactions among cINs of the same age (Southwell et al., 2012). Given the key role Pcdhγc4 plays in the survival of cINs and the fact that Pcdhγc4 is a cell-adhesion molecule that is involved in homophilic cell-cell recognition (Thu et al., 2014), initial cell-cell interactions within the cIN population could be mediated through Pcdhγc4. Other studies have shown that programmed cell death is also regulated at least in part through activity-dependent mechanisms (Duan et al., 2020; Wong et al., 2018, 2022). It remains undetermined what role, if any, Pcdhγc4 plays in the regulation of activity during the peak of programmed cell death. In this regard, it would be interesting to determine whether network activity is perturbed in cINs lacking Pcdhγc4. In addition, loss of vesicular GABA release from cINs and the resulting reduction in inhibition leads to their increased participation in network activity and concomitantly to increased cIN survival (Duan et al., 2020). Hence it would also be interesting to remove the function of vesicular GABA release and of Pcdhγ in cINs and determine their survival during programmed cell death.

Appropriate numbers of cINs are considered essential in the modulation of cortical function. This is ultimately adjusted by a period of programmed cell death once the young cINs have migrated into the cortical plate and have begun to make synaptic connections. Here, we identify the Pcdhγc4 isoform as an essential molecular component in the regulation of cIN programmed cell death. It is likely that some of the primary roles that Pcdhγc4 has in the adjustment of cIN cell number to match the number of excitatory neurons apply also to the human cerebral cortex. Recent work suggests that Pcdhγc4 plays a critical role in human brain development as mutations in this gene are associated with a neurodevelopmental syndrome with progressive microcephaly and seizures (Iqbal et al., 2021). An understanding of the cell-cell interactions that use Pcdhγc4 to regulate cIN cell death should give fundamental insights into how the cerebral cortex forms and evolves.

## Methods

### Animals

All protocols and procedures followed the University of California, San Francisco (UCSF) guidelines and were approved by the UCSF Institutional Animal Care Committee. The following breeders were purchased from the Jackson Laboratory: Ai6, Ai14, Gad1-GFP, *BAC-Nkx2.1-Cre* (*Nkx2.1^Cre^*), and WT C57BL/6J. Pcdhγ-fcon3 (FCON3) mice were obtained from Joshua Sanes at Harvard University. *Pcdhγ^1R1/1R1^(1R1), Pcdhγ^C4KO/C4KO^(C4KO) and Pcdhγ^3R2/3R2^(3R2)* mice were transferred from the Weiner laboratory at the University of Iowa and re-derived, and bred at UCSF.

*Pcdhγ* loss of function mice were obtained by crossing *Pcdhγ^fcon3/fcon3^* mice with *Nkx2.1^Cre^-Ai14-Pcdhγ^fcon3/+^. Pcdhγ^1R1/1R1^(1R1*) and *Pcdhγ^C4KO/C4KO^(C4KO) mice were crossed to Nkx2.1^Cre^-Ai14 breeders to label the Nkx2.1 progenitor cells*.

For cell transplantation experiments, GFP-expressing cIN in embryos was produced by crossing WT C57BL/6J to heterozygous mice expressing green fluorescent protein-expressing (GFP) driven by Gad1 or by crossing Nkx2.1^Cre^;Ai6 breeder to WT C57BL/6J mice. For all, tdTomato-expressing cells derived from MGE embryonic dissections we crossed the various mutant alleles to the *Nkx2.1^Cre^-Ai14 line. Pcdh-γ^fcon3/fcon3^* mutant embryonic tissue was obtained from embryos produced by crossing *Nkx2.1^Cre^-Ai14-Pcdhγ^fcon3/+^* mice with *Pcdhγ^fcon3/fcon3^* mice. *Pcdhγ^1R1/1R1^* mutant tissue was obtained from embryos produced by crossing *Nkx2.1^Cre^-Ai14-Pcdhγ^1R1/1R1^* breeders. *Pcdhγ^C4KO/C4KO^* mutant tissue was obtained from embryos produced by crossing *Nkx2.1^Cre^-Ai14-Pcdhγ^C4KO/+^* breeders. *Gad1-GFP, Nkx2.1^Cre^; Ai6* and *Nkx2.1^Cre^-Ai14* offspring were genotyped under an epifluorescence microscope (Leica), and PCR genotyping was used to screen for *Pcdhγ^fcon3/fcon3^, Pcdhγ^1R1/1R1^ and Pcdhγ^C4KO/C4KO^* alleles in tdTomato positive embryos or reporter negative embryos. All cell transplantation experiments were performed using wild type C57Bl/6 recipient mice P2 to P8. All mice were housed under identical conditions.

### Plasmids

The following plasmids were used in this work: pLenti-CAG-EGFP, pLenti-CAG-Pcd*γ*c4-EGFP.

### Timed pregnant mice

In experiments requiring timed pregnant mice, the day when the sperm plug was observed was considered E0.5. Males were paired with females the night before and checked for plugs early the following day. MGE cells for transplantation were dissected from the fetal forebrain between E12.5 - E15.5 embryos as previously described (Southwell et al., 2012).

### Transplantation

For co-transplantation experiments, the concentration of cells of each genotype was determined using a hemocytometer; the GFP or tdTomato labeled cells were then mixed in similar proportions. To prepare the cells for transplantation, the cells suspension was concentrated by spinning in a table centrifuge for five minutes at 800g(rcf). The supernatant was removed and the resulting pellet was resuspended and mixed in a final volume of 1-6μL of Leibovitz L-15 medium (L15). This concentrated cell suspension was loaded into beveled glass micropipettes (~60-100 μm diameter, Wiretrol 5 μL, Drummond Scientific Company) prefilled with mineral oil and mounted on a microinjector as previously described (Wichterle et al., 1999). The viability and concentration of the cells in the glass micropipette was determined by using 100nl of cells diluted 200X in 10μL of L15 medium and 10μL trypan blue. The number of cells was then determined using a hemocytometer. Cells were injected into neonate mice P2 to P8. Prior to the injection of cells, the recipient mice (C57Bl/6) were anesthetized by hypothermia (~3-5 minutes) and positioned in a clay head mold that stabilizes the skull (Merkle et al., 2007). Micropipettes were positioned at an angle of 0 or 30 degrees from vertical in a stereotactic injection apparatus, and injected between 4-6mm A/P and 0.5 to1.5mm M/L and 0.8 to 0.5 mm in the Z direction. After the injections were completed, the transplanted mice were placed on a warm surface to recover from hypothermia. The mice were then returned to their mothers. Transplantation of cells involving the *Pcdhγ^C4KO/C4KO^* alleles was performed utilizing cells from MGEs that had been cryopreserved (Rodríguez-Martínez et al., 2017).

### Tissue dissection and cell dissociation

MGEs were dissected from E12.5 to E15.5 embryos as previously described (Southwell et al., 2012). Dissections were performed in ice-cold L15 medium and the dissected MGEs were kept in this medium at 4°C. After collecting and genotyping the embryos, MGE with the same genotype were pulled together and mechanically dissociated into a single cell suspension by repeated pipetting in L15 medium containing DNAse I (180μg/ml)using a P1000 pipette. For experiments involving cells from cryopreserved MGEs, cryovials were removed from −80°C, thawed at 37°C for 5 minutes and the content of each tube was resuspended in a 15mL falcon tube containing L15 medium at room temperature. The MGE were washed twice with L15 to remove residual DMSO and dissociated as above.

### Cryopreservation of MGE in toto

Dissected MGEs from each embryo were collected in 500 μL L15 medium and kept on ice until cryopreserved. MGEs were resuspended in 10% DMSO in L15 medium and cryopreserved as previously described (Rodríguez-Martínez et al., 2017). Vials were cooled to −80°C at a rate of −1°C/minute in a Nalgene™ Mr. Frosty Freezing Container and transferred the next day to liquid nitrogen for long term storage. Importantly, tissue for genotyping was collected from each embryo and labeled with a code name matching the codename of each cryotube used for the cryopreservation.

### Cell counting

GFP positive cells and tdTomato-positive cells were counted in all layers of the neocortex. Cells that did not display neuronal morphology were excluded from all quantifications. The vast majority of cells transplanted from the E13.5 MGE exhibited neuronal morphologies in the recipient brain. Cells at the injection site were not counted. Most cells migrated away from the injection site, but few remained trapped. Cells outside the cortex were not included in the quantification. In some injections, cells migrated to other parts of the brain, including the hippocampus and striatum. Images of co-transplanted cells were acquired on a SP8 (Leica) confocal microscope with a 10X magnification. Images of transplanted cells infected with lentivirus expressing the Pcdhγc4 were acquired on an SP8 confocal with a 10X magnification and digital zoom=2x. Only the region of the cortex containing GFP or tdTomato-positive cells was imaged. GFP and tdTomato-positive cells were counted using Neurolucida (MBF). For all experiments, transplanted cells were counted from coronal sections along the rostral-caudal axis and in at least 10 sections were counted per animal; only sections containing more than 10 cells per condition were used. Data is presented as the fraction of transplant-derived cells that express GFP and/or tdTomato. This fraction does not reflect the absolute number of cells, but their relative contribution to the overall population of transplant-derived cells at different DAT. For experiments involving the expression of lentiviral driven GFP or *Pcdhγ* in transplanted cINs, survival was determined by comparing the fraction of infected (tdTomato+GFP+) cells to the fraction of noninfected cells (tdTomato+GFP-) from the total number of transplanted cells (tdTomato+) in each brain section.

### Viral vector subcloning

All lentiviral plasmids were cloned from the backbone construct pLenti-CAG-ires-EGFP (addgene plasmid #122953). A Kozak sequence was added at the start of the coding region for all genes cloned. Plasmids were cloned using the Gibson Cloning Kit (NEB). The pLenti-CAG-EGFP construct was cloned by removing the ires sequence in between the BxtXI and BamHI restriction sites from the backbone construct. To clone the pLenti-CAG-Pcdhγc4-EGFP (fusion construct), the pLenti-CAG-ires-EGFP plasmid was digested with BamHI and BstXI to remove the IRES sequence. A PCR generated *Pcdhγc4* coding sequence lacking the stop codon and containing a two amino acid (Ser, Arg) linker was cloned upstream of the GFP coding sequence. This construct was used to express Pcdhγc4 fused to GFP.

### Viruses

All lentivirus used in this study were made in the laboratory. Briefly viruses were produced in Lenti-X 293T cells (TakaraBio). Cells were grown to ≥90% confluency in maintenance media (DMEM/F-12, 15 mM HEPES, 2.5 mM GlutaMAX, 1% Pen/Strep, 10% FBS) in 15cm plates coated with Poly-D-lysine (Sigma-Aldrich P7405) at final concentration of 0.1mg per mL. Once cells reached the desired confluency, media was changed to DMEM/F-12 + 2% FBS. Cells were transfected using *Trans*IT®-293 Transfection Reagent (Mirusbio) and Opti-MEM (Thermofisher). After 6 to 12 hours post-transfection, the media was changed to lentivirus production media (DMEM/F-12, 15 mM HEPES, 2.5 mM GlutaMAX, 1% Pen/Strep, 2% FBS) and 60μLof ViralBoost Reagent was added (ALSTEM) per 15cm plate. Virus supernatant was collected 48 hour post-transfection, filtered through a 45μm filter, and precipitated with Lentivirus Precipitation Solution (ALSTEM) overnight following manufacturer’s instructions. The viral pellet from two 15cm plates was concentrated in a final volume of 100μl for Pcdh constructs and 200μl for the control construct.

### Viral infection of MGE precursor cells

Following the dissociation of the MGEs, cells were concentrated by spinning in a table centrifuge for five minutes at 800g. The cell pellet was subsequently resuspended in an equal volume of Lentivirus and L15 medium. The cell-virus mix was incubated at 22-32°C at 1000rpm (190rcf) for 3 hours, mixing every 30 minutes.

### Immunostaining

P21 and older mice were fixed by transcardiac perfusion with 10mL of ice-cold PBS followed by 10mL of 4% formaldehyde/PBS solution; transcardiac perfusion of 5ml of either solution was used for P15 and younger mice. Brains were incubated overnight in the same fixative (12-24hrs) at 4°C, then rinsed with PBS and cryoprotected in 30% sucrose/PBS solution for 48 hours at 4°C. Unless otherwise stated, immunohistochemistry was performed on 30 μm floating sections in Tris Buffered Saline (TBS) solution containing 10% normal donkey serum, 0.5% Triton X-100 for all procedures on postnatal mice. All washing steps were done in 0.1% Triton X-100 TBS for all procedures. Sections were incubated overnight at 4°C with selected antibodies, followed by incubation at 4°C overnight in donkey secondary antibodies (Jackson ImmunoResearch Laboratories). Brain sections that had been transplanted with lentivirus infected MGE cells were incubated two nights in primary antibodies and overnight with secondary antibodies to enhance the viral expressed reporter GFP. For cell counting and *post hoc* examination of marker expression, sections were stained using chicken anti-GFP (1:2500, Aves Labs, GFP-1020, RRID:AB_10000240), rabbit anti-RFP (Rockland), rat anti-tdTomato (Kerafast). For staining with the Pcdhγc5 antibody, tissue fixation was performed as stated above. Sixteen um sections of cortical tissue were dried directly onto paraffin-subbed slides before being washed 3X for 5 min. each in TBS. Sections were incubated in a blocking solution with 2.5% BSA, 0.5% TritonX, in TBS for 2 hours at room temp. Slides were removed from the blocking buffer and washed again 3X for 5 min. Primary antibodies were diluted in 2.5% BSA 0.25% TritonX 0.02% in TBS overnight at 4°C. Antibodies used; Pcdh*γ*c5 (Invitrogen, 1:150, RRID:AB_2724958), Ctip2 (Abcam, 1:200, PRID:AB_2064130), Cux1 (Santa Cruz Biotechnology, 1:100 AB_2261231), GABA (Sigma-Aldrich,1:250, RRID:AB_477652). Sections were then washed 3X for 5 min. before being incubated with host-appropriate secondaries for 2 hours at room temp at a 1:500 dilution in.

### RNAScope

Brains of *Nkx2.1^Cre^; Ai6* animals at multiple ages (P5, P7, and P14) were fixed, postfixed, and cryoprotected (as described in *Immunostaining*). Brains were cryosectioned and mounted directly onto microscope slides before being stored at −80 °C. The RNA scope experiments were completed as per the manufacturer’s instructions (RNAscope multiplex fluorescent reagent kit v2, Advanced Cell Diagnostics, Inc.). Sections were equilibrated to room temperature by one 10 min incubation at −20°C before a 10 min incubation at room temperature. Brain sections were dehydrated using increasing concentrations of Ethanol (50%, 70% and 100%) for 5 min incubations at room-temperature. Sections were then incubated with RNAscope Hydrogen Peroxide solution for 10 min. at room temperature, before 4 washes in distilled water. To perform the target retrieval step of the protocol, slides were incubated in distilled water for 10 seconds and then RNAscope 1X target retrieval buffer for 15 min. using an Oyster brand steamer at ~99°C. Sections were then washed for 15 seconds in distilled water before being transferred to 100% ethanol for 3 min. at room-temperature. Slides were then dried in an incubator at 60°C for 5 min. In order to reduce reagent usage, a hydrophobic barrier pen was used to surround the tissue. After the barrier was dry, tissue was incubated in ~5 drops of RNAscope Protease III, and incubated at 40°C for 30 min. using a HybEZ Humidity Control Tray and HybEZ oven.

Tissue was then incubated in the *Pcdhγc4* (probe 835791-C3) and *Pcdhγc5* (probe 1118501-C2) probes for 2 hours at 40°C. After incubation, Probes were hybridized with AMP1, AMP2, and AMP3 (in that order) through individual incubations at 40°C for 30 min (AMP3 incubation 15 min. incubation). Our probes were Channel-2 (*Pcdhγc5*) and Channel 3 (*Pcdhγc4*), so we skipped development HRP-C1 signal and continued directly to the development of the HRP-C2 stepRNAscope Multiplex FL v2 HRP-C2. Tissue was incubated with 3-4 drops of RNAscope multiplex v2 HRP-C2 solution for 15 min at 40°C, followed by a 2 min wash in 1X RNA scope wash buffer. Then, tissue was incubated with Cy3 (Thermo Fisher Scientific, 1:1000) diluted in TSA buffer (Advanced Cell Diagnostics, Inc.), followed by another 2 min wash in 1X Wash Buffer. This process was completed again using the RNAscope multiplex v2 HRP-C3 solution and Cy5 (Thermo Fisher Scientific, 1:1000). Slides were washed 3X in distilled water for 5 min., before being mounted using Fluoromount-G, DAPI mounting media.

### Analysis of Previously Published Single Cell RNA Sequencing Datasets

Data from Tasic et al., 2018 was obtained via the Broad Institute Single Cell Portal (https://singlecell.broadinstitute.org/single_cell/study/SCP6/a-transcriptomic-taxonomy-of-adult-mouse-visual-cortex-visp). Cells were grouped by GABAergic vs Glutamaterigc designation or cluster-type, as determined by the data generators. RPMK values for each cell in each cluster-type were averaged and plotted using PRISM. Data obtained from Yao et al., 2021 was obtained from the UCSC Cell Browser (https://cells.ucsc.edu/?ds=allen-celltypes+mouse-cortex+mouse-cortex-2019). Cells cells were grouped based on either GABAergic or Glutamatergic designations or by cluster-type determined by the data generators. Absolute transcript level Absolute transcript values for each cell in each cluster-type were averaged and plotted using PRISM.

### Statistical Analysis

Quantifications were performed by two different people, one of which was blinded to the genotype. For transplantation experiments, we estimated the average mutant and WT survival ratios in each mouse by counting the transplanted mutant and wild-type cells in each of ~10 sections. We used an ANOVA to determine whether the survival ratios in 6DAT mice were significantly different from those in 21DAT mice.

---

**Figure 2 Supplement.**
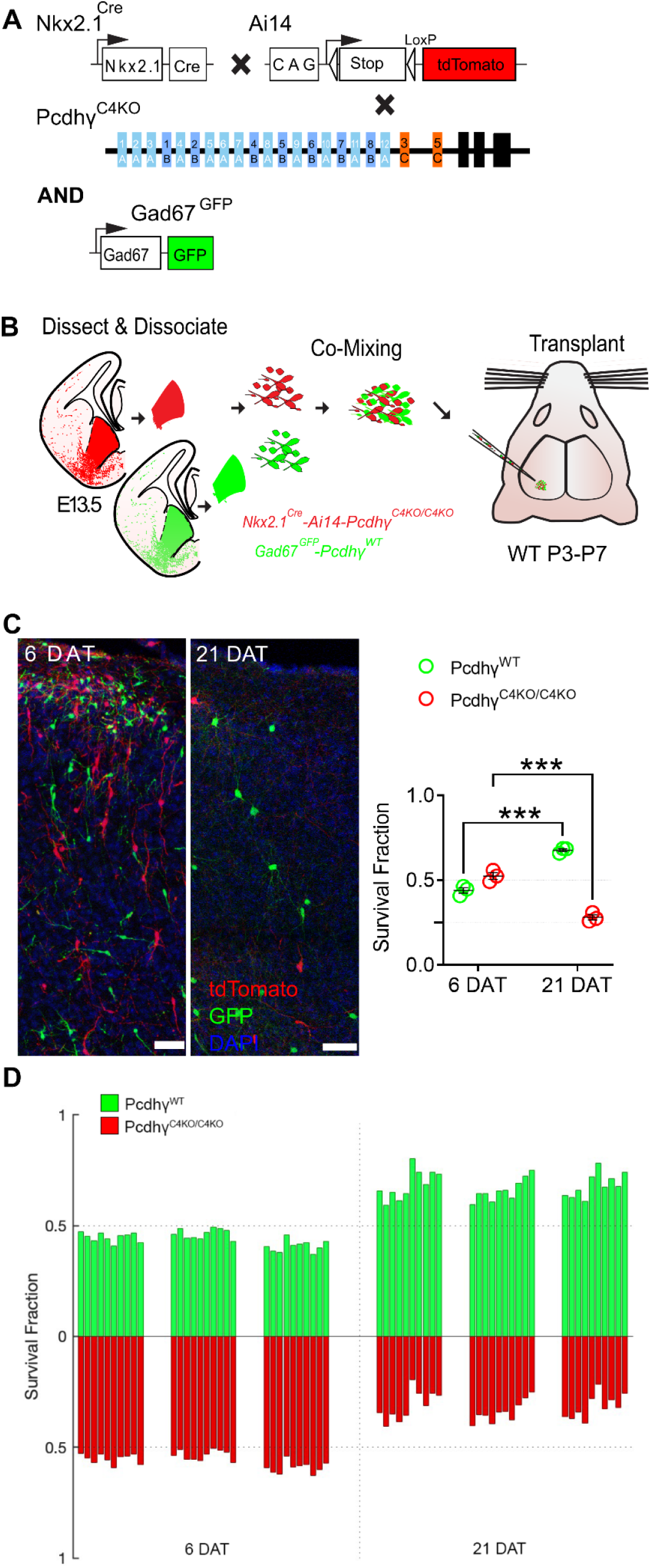
Genetic deletion of Pcdhγc4 increased cell death in MGE-derived cINs. **A.** Diagram of genetic crosses. Pcdhγ^C4KO^ homozygous MGE cells were labeled via the MGE-specific genetic reporter *Nkx2.1^Cre^* mice that also carry the conditional Ai14 allele. Control Pcdhγ^WT^ MGE cells were labeled with GFP via the Gad67-GFP report mice. **B.** Schematics of transplantation protocol. The MGEs of Pcdhγ^C4KO^ homozygous mutant or control E13.5 embryos were dissected, dissociated, and mixed in similar proportions. The mixture of GFP+ (Pcdhγ-WT) and tdTomato+ (Pcdhγ^C4KO/C4KO^) cells were grafted into the cortex of WT neonate mice. **C.** Left - Confocal images from the cortex of 6 and 21 DAT mice. The transplanted cells were labeled with GFP (Pcdhγ^WT^) or tdTomato (Pcdhγ^C4KO/C4KO^). Right - Quantifications (shown as survival fraction) of surviving GFP or tdTomato labeled MGE-derived cINs at 6 and 21 DAT. Both the GFP and tdTomato labeled cells undergo programmed cell death between 6 and 21DAT, but Pcdhγ^C4KO/C4KO^ cells are eliminated at higher rates. **D.** Survival fraction quantification from (C) shown by the brain section (each bar) and separated by animals at 6 and 21 DAT. Scale bar = 50 um, Nested-ANOVA, ***p = 0.0002, n = 3 mice per time point and 10 brain sections quantified per mouse.

**Figure 3 - supplement.**
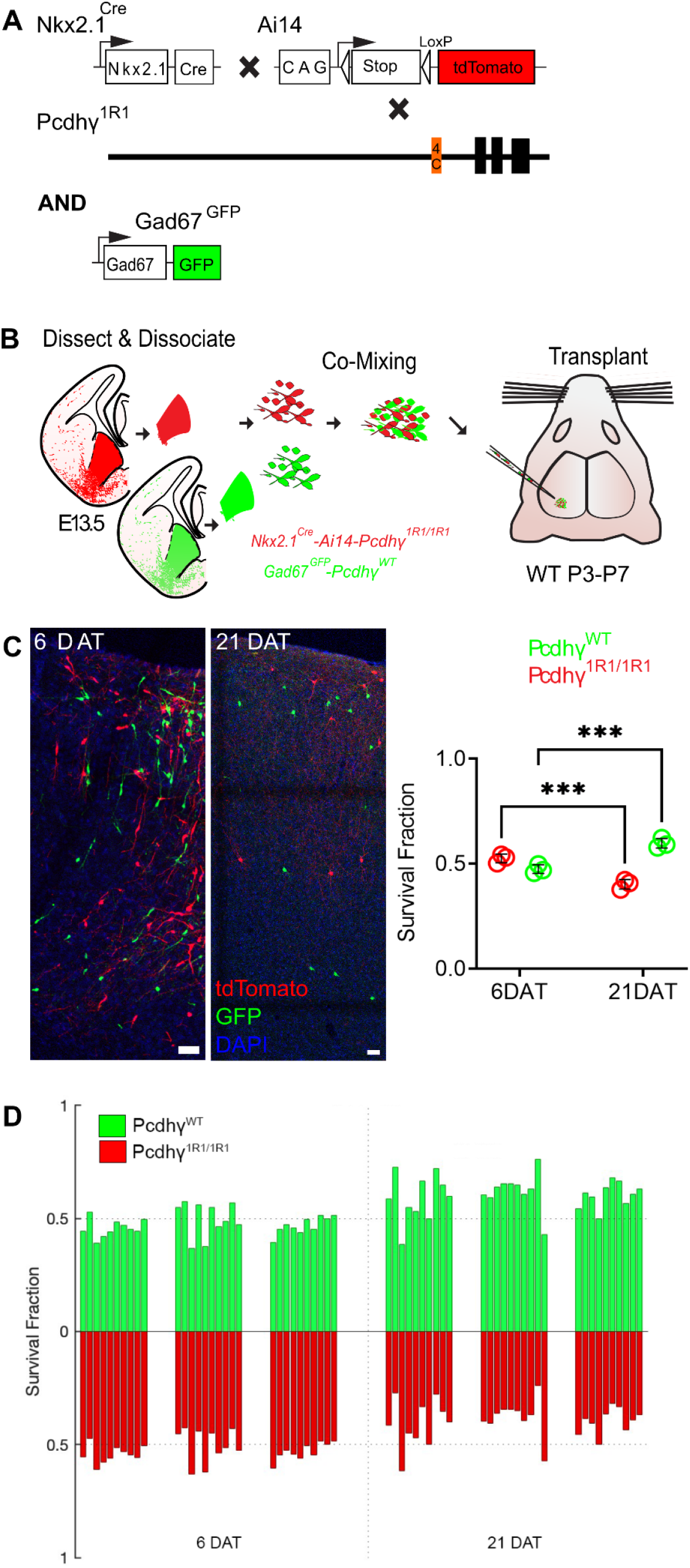
Expression of Pcdhγc4 is sufficient for the survival of most MGE-derived cINs. **A.** Diagram of genetic crosses between MGE/POA-specific reporter *Nkx2.1^Cre^*;Ai14 and Pcdhγ^1R1^ mice. Control cells were obtained from *Gad67-GFP* embryos. **B.** Schematics of transplantation protocol. The MGEs from E13.5 Pcdhγ^1R1^ homozygous or control embryos were dissected, dissociated, and mixed in similar proportions. The mixture of GFP+ (Pcdhγ^WT^) and tdTomato+ (Pcdhγ^1R1/1R1^) cells was grafted into the cortex of WT neonate mice. **C.** Left - Confocal images from the cortex of 6 and 21 DAT mice. The transplanted cells are labeled with GFP (Pcdhγ^WT^) or tdTomato (Pcdhγ^1R1/1R1^). Right - Quantifications (shown as survival fraction) of surviving MGE-derived cINs at 6 and 21 DAT. Both the transplanted GFP and tdTomato-labeled cells undergo programmed cell death between 6 and 21 DAT, but the Pcdhγ^1R1/1R1^ cells are eliminated at higher rates. **D.** Survival fraction quantification from (C) shown by the brain section (each bar) and separated by animals at 6 and 21 DAT. Scale bar = 50 um, Nested-ANOVA, ***p = 0.009, n = 3 mice per time point and 10 brain sections per mouse

**Figure 4 supplement.**
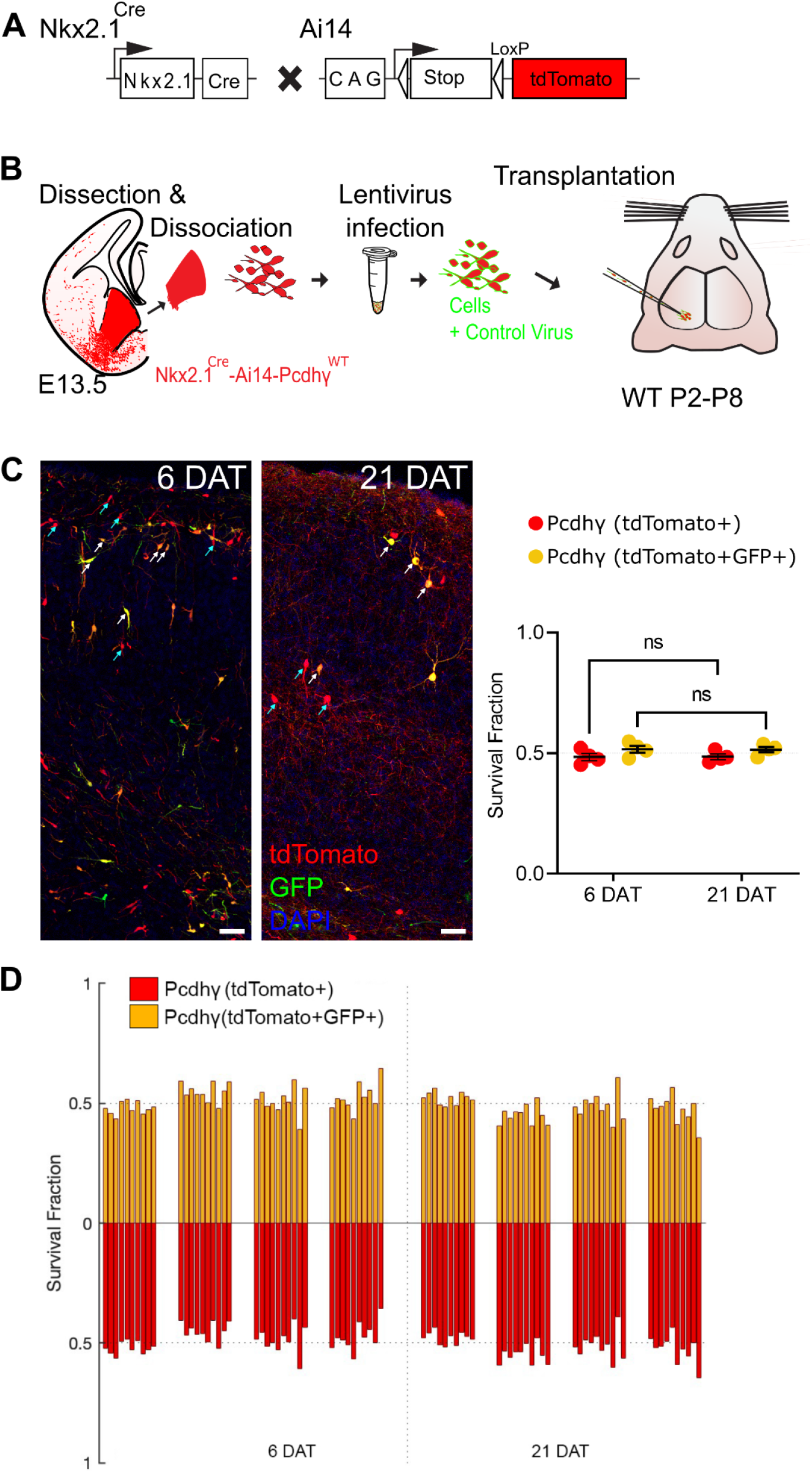
Infection with lentivirus and expression of GFP does not affect the survival of cINs. **A** Diagram of mouse crosses to obtain Pcdhγ^WT^ MGE cells labeled with MGE-specific genetic reporter. **B.** Schematic of lentiviral infection and transplantation of MGE cIN precursors. The MGEs of *Nkx2.1^Cre^-Ai14* embryos that carry Pcdhγ^WT^ were dissected, dissociated, and infected in suspension with lentivirus expressing GFP. The infected cells were grafted into the cortex of WT recipient mice P0-P8. **C.** Confocal acquired images of the transplanted cINs in the cortex at 6 and 21 DAT. Notice that all transplanted cells are labeled with tdTomato (red), while cells expressing virally-driven GFP are labeled in green. Quantifications of the tdTomato+ only (teal arrows) or tdTomato+GFP+ (yellow cells, white arrows) cells are shown as the fraction of cells from the total tdTomato+ cells at 6 and 21 DAT. **D.** Survival fraction quantification from (C) shown by the brain section (each bar) and separated by animals at 6 and 21 DAT. Scale bar = 50 um, Nested-ANOVA, ns = not significant, n = 4 mice per time point and 10 brain sections per mouse from one transplant cohort.

## Author contributions

Conceptualization, W.R.M, D.M.S, B.R., M.P.S., A.R.H., J.A.W., B.R.; Methodology, W.R.M., D.M.S.; Investigation, W.R.M, D.M.S., F.D.; Technical Assistance, F.D., R.R., A,B.; Writing – Original Draft, W.R.M, A.A.B., D.M.S.; Writing – Review & Editing, A.A.B., W.R.M, D.M.S, M.P.S., A.R.H., J.A.W., B.R.,; Funding Acquisition, A.R.H., M.P.S., J.A.W., A.A.B.,; Supervision, A.R.H., M.P.S., J.A.W., A.A.B.

## Acknowledgements

This work was supported by NIH Grant R01MH122478 to A.A.B., A.R.H., and M.P.S.; NIH Grants R01 NS028478 and R01 EY02517; and a generous gift from the John G Bowes Research Fund to A.A.B.; National Institutes of Health Grants R01 NS055272 and R21 NS090030 to J.A.W.; National Institutes of Health Grants R01NS116598; Hearing Research Inc.; the PBBR Breakthrough Fund; and the Coleman Memorial Fund. to A.R.H.; NIH Grants R01EY002874 and R01EY031059 to M.P.S. A.A.B. is the Heather and Melanie Muss Endowed Chair and Professor of Neurological Surgery at UCSF. M.P.S. is a recipient of the Research to Prevent Blindness Disney Award for Amblyopia Research.

